# Lysosomal cystine mobilization shapes the response of mTORC1 and tissue growth to fasting

**DOI:** 10.1101/606541

**Authors:** Patrick Jouandin, Zvonimir Marelja, Andrey A Parkhitko, Miriam Dambowsky, John M Asara, Ivan Nemazanyy, Christian C. Dibble, Matias Simons, Norbert Perrimon

## Abstract

Adaptation to nutrient scarcity involves an orchestrated response of metabolic and signaling pathways to maintain homeostasis. We provide evidence that lysosomal export of cystine coordinates remobilization of internal nutrient stores with reactivation of the growth regulator TORC1 signaling upon fasting in the *Drosophila* fat body. Mechanistically, cystine is reduced to cysteine and metabolized to acetyl-CoA by consuming lipids. In turn, acetyl-CoA retains carbons from alternative amino acids in the form of TCA cycle intermediates, thereby restricting amino acids availability, notably aspartate. This process limits TORC1 reactivation to maintain autophagy and allows animals to cope with starvation periods. We propose that cysteine metabolism mediates a communication between lysosomes and mitochondria to maintain the balance between nutrient supply and consumption under starvation, highlighting how changes in diet divert the fate of an amino acid into a growth suppressive program.

**One Sentence Summary:** Lysosomal cysteine recycling is a metabolic break that maintains autophagy upon starvation through coenzyme A synthesis.

## Main Text

Organisms constantly cope with variations in diet by adjusting metabolism, and specific organs impact systemic nutrient homeostasis. Upon fasting, the liver remobilizes nutrients through gluconeogenesis and β-oxidation of fatty acids to support peripheral tissue function (*1, 2*). Variation in nutrient availability induces parallel changes in signaling pathways activity, which resets intracellular metabolic turnover. The TORC1 (Target of Rapamycin Complex 1) signaling pathway integrates sensing of amino acids and other nutrients with signals from hormones and growth factors to promote growth and anabolism (*3*). Nutrient scarcity inhibits TORC1 to limit growth and promote catabolic programs, including autophagy that recycles internal nutrient stores to promote survival (*4*). Autophagy involves the sequestration of cytosolic material into autophagosomes that fuse with lysosomes for cargo degradation and recycling. The lysosomal surface is also the site where nutrient and growth factors sensing pathways converge to activate TORC1. Degradation within lysosomes generates new amino acids that in turn fuel metabolic pathways, including the tricarboxylic acid (TCA) cycle and gluconeogenesis, and reactivate TORC1, altogether maintaining minimal anabolism and growth (*5-8*). However, how organisms regulate the limited pools of remobilized nutrients and balance homeostatic metabolism with anabolic TORC1 activity over the course of starvation is poorly understood.

## Results

To study the regulation of metabolism and TORC1 signaling *in vivo*, we used the *Drosophila* larval fat body, an organ analogous to the liver and adipose tissue in mammals. The fat body responds to variation in nutrient availability through TORC1 signaling, which in turn regulates systemic larval growth rate through the secretion of growth factors from distant organs (*9, 10*). When larvae were completely deprived of their food source (but kept otherwise hydrated), TORC1 signaling in the fat body was acutely downregulated at the onset of starvation, as expected. However, prolonged starvation led to partial and progressive reactivation of TORC1 over time (Fig. 1A). This process was dependent on autophagy induction (Fig. S1A, B), consistent with autophagy regulating amino acid recycling and TORC1 reactivation in mammalian cells (*6, 8*). To understand how changes in nutrients and metabolites intersect with mTORC1 signaling during this fasting response, we initially took two complementary unbiased approaches: 1) a targeted mass-spectrometry based screen for polar metabolites altered during starvation and 2) a larval growth screen to test the effects of individual amino acids during starvation.

**Figure 1:**
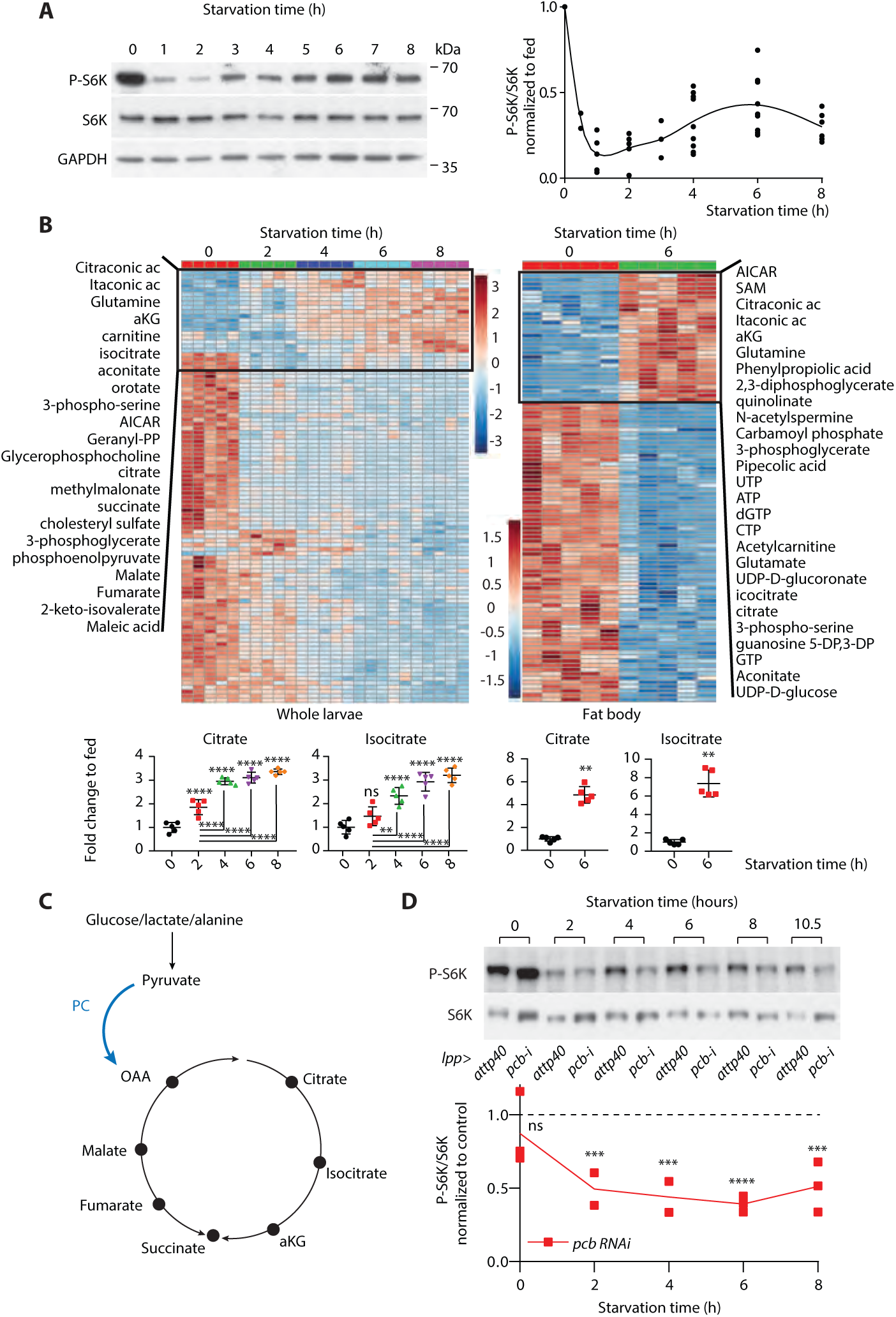
TORC1 reactivation upon prolonged fasting correlates with increase of TCA cycle intermediates. A) Prolonged fasting leads to TORC1 reactivation. Phosphorylation levels of the direct TORC1 target S6K in dissected fat bodies from larvae starved on PBS. B) Heat map metabolite levels (LC-MS/MS) from whole mid-third instar larvae (left) or dissected fat bodies (right), fed (0h) and starved on PBS. Lower panel, individual plots from the same dataset. C) Schematic of anaplerosis through pyruvate carboxylase PC. D) Knockdown of *pcb* (PC) suppresses mTORC1 reactivation upon prolonged fasting. B, D: Mean +/- SD, ^ns^, P≥0.05; *, P≤0.05; **, P≤0.01; ***, P≤0.005; ****p≤0.0001 (See methods for details).

Metabolic profiling of whole animals in matching conditions revealed depletion of most metabolites over the course of starvation (Fig. 1B). By contrast, TCA cycle intermediates accumulated over time, in particular citrate/ isocitrate. This was not due to defective TCA cycle activity upon fasting evidenced by functional U-^13^C_6_-glucose oxidation in the TCA cycle and NADH/NAD^+^ ratios at equilibrium between fed and fast conditions (Fig. S1C, D). Similar metabolomics profiles were observed in fat bodies dissected out from fed and starved animals (Fig. 1B), indicating that TORC1 reactivation in the fat body correlates with accumulation of TCA cycle intermediates, in particular citrate / isocitrate. To test a link between changes in the TCA cycle and TORC1 reactivation, we targeted pyruvate carboxylase (PC). PC serves an anaplerotic function by replenishing TCA cycle intermediates through oxaloacetate (Fig. 1C), a process that regulates gluconeogenesis and regeneration of amino acids (*1*). Knockdown of PC (*pcb/CG1516*) in the larval fat body elevated alanine levels (Fig. S1E), consistent with alanine as an important PC substrate (*1*). However, it suppressed TORC1 reactivation upon fasting (Fig. 1D), suggesting a functional link between the TCA cycle and TORC1 signaling upon fasting.

In our second screen, we tested the effects of single amino acid supplementation on larval growth upon fasting (see Supplementary Materials). Unexpectedly, this screen revealed cysteine as a unique amino acid associated with growth suppressive properties (Text S1; Fig. 2A; Fig. S2A-D). Again, this process involved changes in TORC1 signaling as cysteine treatment partially inhibited TORC1 activity in fat bodies and constitutive activation of TORC1, in the absence of the TORC1 regulator Gator1 (*nprl2*^*-/-*^) (*11*), partially restored growth under cysteine supplementation (Fig. S2E, F).

**Figure 2:**
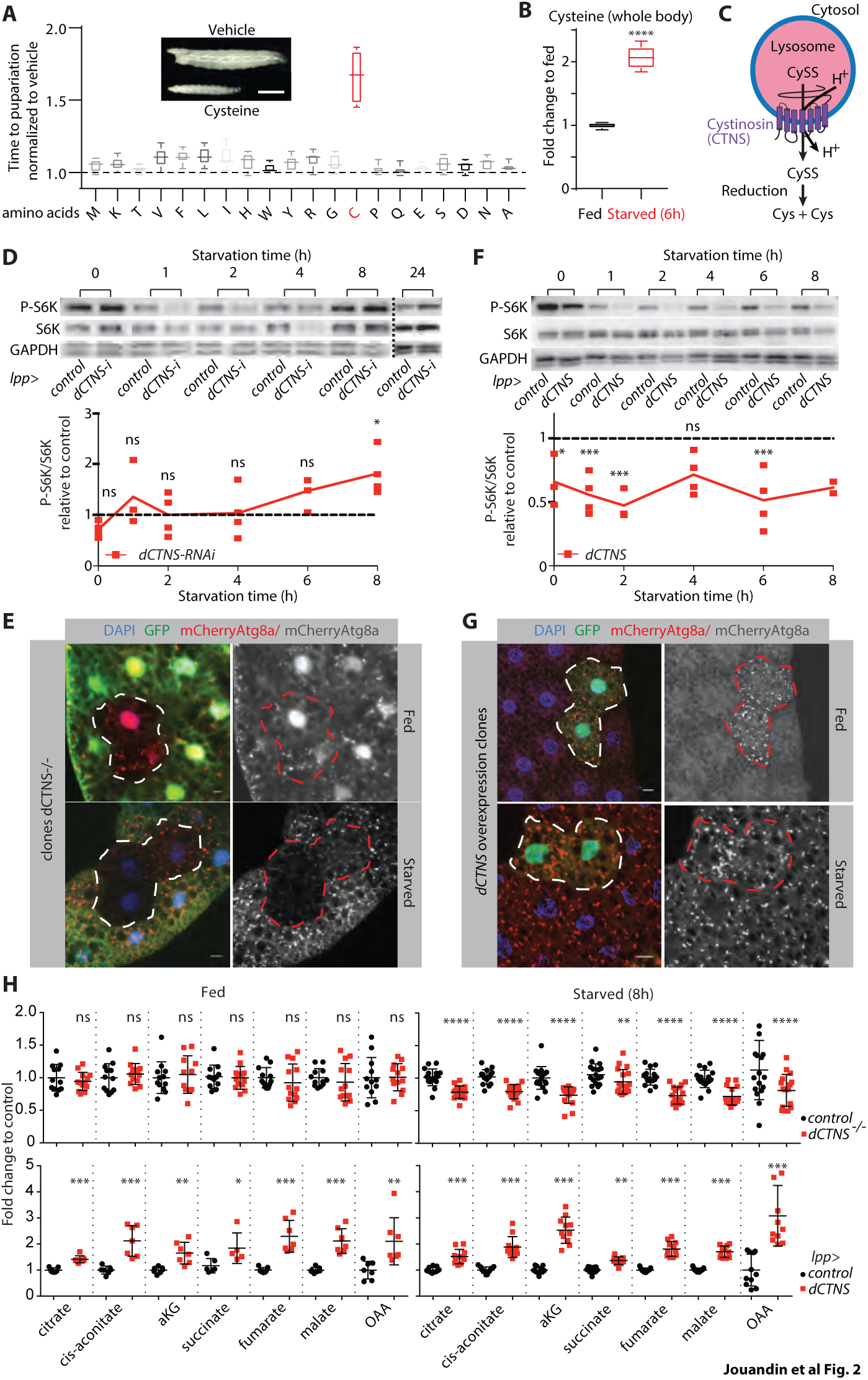
Lysosome-derived cysteine promotes elevation of TCA cycle intermediates and antagonizes TORC1 reactivation. A) Amino acid screen reveals cysteine as a growth suppressor. Time to pupariation for larvae fed a minimal diet supplemented with the indicated amino acids all along development (see supplementary methods for concentration. Cysteine is 5 mM). B) Cysteine levels in control whole control larvae (*lpp>attp40*). N=5. C) Schematic of lysosomal cystine (CySS) efflux through cystinosin/ *CTNS*. D) Loss of cystinosin (*lpp>dCTNS RNAi)* leads to higher TORC1 reactivation upon prolonged starvation. P-S6K levels in dissected fat bodies from larvae transferred to a fast diet. *w-i*, control background E) Cystinosin maintains autophagy during fasting. *dCTNS*^*-/-*^ clones (non-GFP, outlined) in 80h AEL (after egg laying) larvae expressing mCherry-Atg8a starved for 8h. Scale bar 10 μm. F) *dCTNS* overexpression suppresses TORC1 activity. P-S6K levels in dissected fat bodies from larvae transferred to a fast diet. GFP-i, control background. G) *dCTNS* overexpression induces autophagy. *dCTNS* overexpression clones (GFP marked, outlined) in 80h AEL larvae. Scale bar 10 μm. H) Cystinosin regulates TCA cycle intermediates levels upon starvation. Relative metabolites levels measured by LC-MS/MS in 80h AEL larvae fed or starved 8h on PBS. Controls are CTNS^+/-^ (upper graphs) or GFP-i (lower graphs). B,D,F,H) Mean +/- SD, ^ns^, P≥0.05; *, P≤0.05; **, P≤0.01; ***, P≤0.005; ****p≤0.0001 (Details in Methods).

The results of our two screens led us to explore the relationship between cysteine metabolism, the TCA cycle, and TORC1 during the fasting response. Importantly, we found that cysteine levels increased upon fasting (Fig. 2B), consistent with previous reports in yeast (*12*), and that cysteine treatment increased the levels of TCA cycle intermediates, particularly upon fasting (Fig. S2G). To further understand this effect of cysteine, we searched for its intracellular source under starvation conditions. Physiologically, cellular cysteine can either be synthesized in the cytosol or transported as cystine from the extracellular space by the Xc^-^ antiporter (*13, 14*) and by cystinosin from the lysosome (*15*) (Fig. 2C). Abolishing autophagy in the fat body decreased cysteine levels upon fasting (Fig. S3A), suggesting that lysosomal function regulates cysteine balance. Thus, we analyzed the role of the lysosomal cystine transporter cystinosin in recycling cysteine upon fasting. Cystinosin, encoded by *CTNS* in mammals, is mutated in the lysosomal storage disorder cystinosis and has previously been implicated in mTORC1 signaling and autophagy (*15-20*). Endogenous tagging of the *Drosophila* ortholog *CG17119* (hereafter referred to as *dCTNS*) confirmed its specific lysosomal localization in fat body cells. *dCTNS*^-/-^ larvae showed accumulation of cystine (Fig. S3B-D), consistent with a role for cystinosin in lysosomal cystine transport. In fed conditions, control and *dCTNS*^*-/-*^ larvae showed similar cysteine levels, likely reflecting dietary intake as the main source of cysteine. By contrast, upon fasting, cysteine levels dropped in *dCTNS* ^*-/-*^ larvae (Fig. S3E). Conversely, *dCTNS* overexpression increased cysteine levels, which caused a developmental delay similar to cysteine treatment (Fig. S3F, G). We therefore concluded that cystinosin recycles cysteine from the lysosome upon fasting and analyzed its effect on TORC1 reactivation and the TCA cycle. While *dCTNS* knockdown in larval fat body did not affect TORC1 inhibition at the onset of starvation, it slightly increased TORC1 reactivation upon prolonged starvation (Fig. 2D). Analysis of *dCTNS*^*-/-*^ fat body clones showed that increased TORC1 signaling was cell-autonomous and sufficient to compromise maintenance of autophagy in starvation. Accordingly, treatment with the TORC1 inhibitor rapamycin was able to restore autophagy in *dCTNS-*deficient cells (Fig. 2E, Fig. S3H, I). Conversely, *dCTNS* overexpression caused downregulation of TORC1 and induced autophagy (Fig. 2F, G). Metabolic profiling of *dCTNS*^*-/-*^ animals showed a depletion of TCA cycle intermediates specifically upon fasting. In contrast, they accumulated following overexpression of *dCTNS* in fed and starved animals (Fig. 2H).

To test the physiological importance of cysteine recycling upon fasting, we analyzed the role of *dCTNS* on starvation resistance. In fed condition, *dCTNS* deficiency in fat body cells neither affected larval development nor the lifespan of adult flies (Fig. 3A, B). However, in fasted animals, *dCTNS* deficiency caused a developmental delay and shortened lifespan (Fig. 3B, C). This starvation sensitivity was rescued by low concentrations of rapamycin, indicating a key role for altered TORC1 signaling and autophagy in mediating *dCTNS* growth phenotypes (Fig. 3D and see text S2). The role of *dCTNS* upon fasting was dependent on its cystine transport function as treatment with either cysteamine (which exports cystine out of the lysosome independently of cystinosin (*21*)) or low concentrations of cysteine (that did not affect development of control animals) rescued the starvation sensitivity of *dCTNS*^*-/-*^ animals (Fig. 3E-G). Altogether, we demonstrate the physiological importance of cysteine recycling from the lysosome upon fasting, which supports the elevation of TCA cycle intermediates and antagonizes TORC1 reactivation to maintain autophagy.

**Figure 3:**
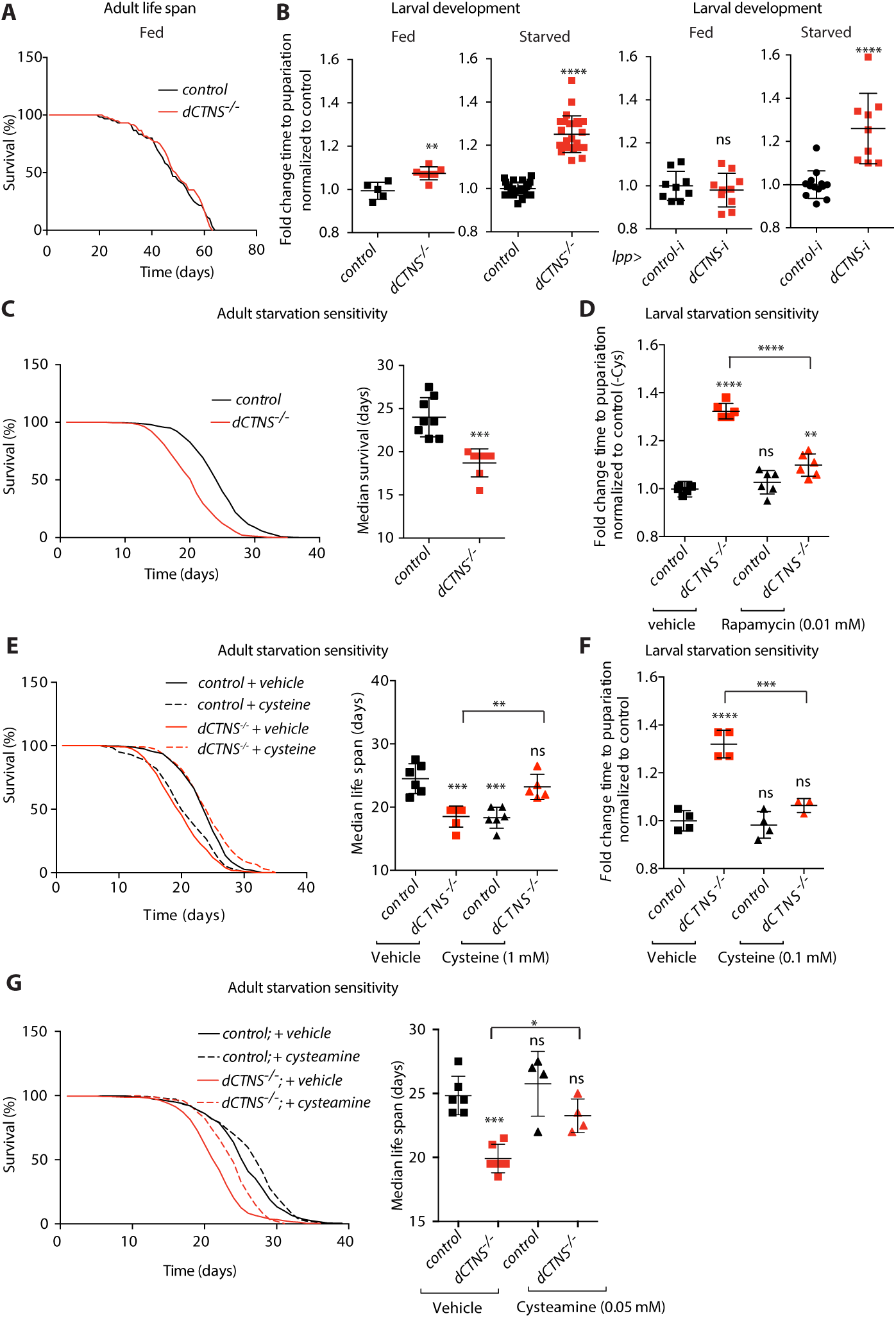
*dCTNS* controls resistance to starvation through cysteine efflux and TORC1. A) Cystinosin does not affect lifespan in fed condition. Lifespan of control (*w*^1118^) and *dCTNS*^*-/-*^ animals fed a standard diet. N=2. B) Cystinosin in the fat body controls starvation resistance during development. Fold change time to pupariation for larvae of indicated genotype grown on fed or fast food. C) Cystinosin controls starvation resistance of adult animals. Lifespan of control (*w*^1118^) and *dCTNS*^*-/-*^ animals fed a chemically defined starvation medium. D-F) Rapamycin and cysteine treatments rescue starvation sensitivity of *dCTNS*^*-/-*^ animals. Developmental time of larvae on starvation food supplemented with 0.01 mM rapamycin (d) or 0.1 mM cysteine (f) and lifespan of adult flies on chemically defined media containing 1 mM cysteine (e). Control is *dCTNS*^*+/-*^ (D, F) and *w*^1118^ (e). G) Cysteamine treatment restores starvation resistance of *dCTNS*^-/-^ animals. Lifespan of control (*w*^1118^) and *dCTNS*^*-/-*^ animals fed a chemically defined starvation medium supplemented with 0.5 mM cysteamine or vehicle. b-g) Mean +/- SEM; ^ns^, P≥0.05; *, P≤0.05; **, P≤0.01; ***, P≤0.005; ****, P≤0.0001 (See statistic details in Methods).

Next, we sought a mechanistic link between cysteine metabolism and the TCA cycle and performed cysteine tracing experiments. For this, we supplemented a fasted diet with either U-^13^C-cysteine or ^13^C_3_ ^15^N_1_-cysteine. Consistent with cysteine metabolism in mammals, this revealed three major cysteine fates: glutathione (GSH), taurine, and coenzyme A (CoA) (Fig. 4A; S4A). The CoA derived from labeled cysteine also contributed significantly to the formation of acetyl-CoA in the fat body (Fig. 4A, B; S4B). Under fasting conditions, the acetyl group of this acetyl-CoA pool is mainly provided through β-oxidation of fatty acids (Fig. 5A). Accordingly, *dCTNS* overexpression upon fasting elevated the level of acetyl-CoA and decreased the level of acyl-carnitines and fatty acids. This phenotype was mirrored in fasted *dCTNS* KO animals (Fig. 5B, C; S5A), suggesting that during fasting, lysosomal cysteine mobilization is rate limiting for *de novo* CoA metabolism and supports acetyl-CoA synthesis through β-oxidation.

**Figure 4:**
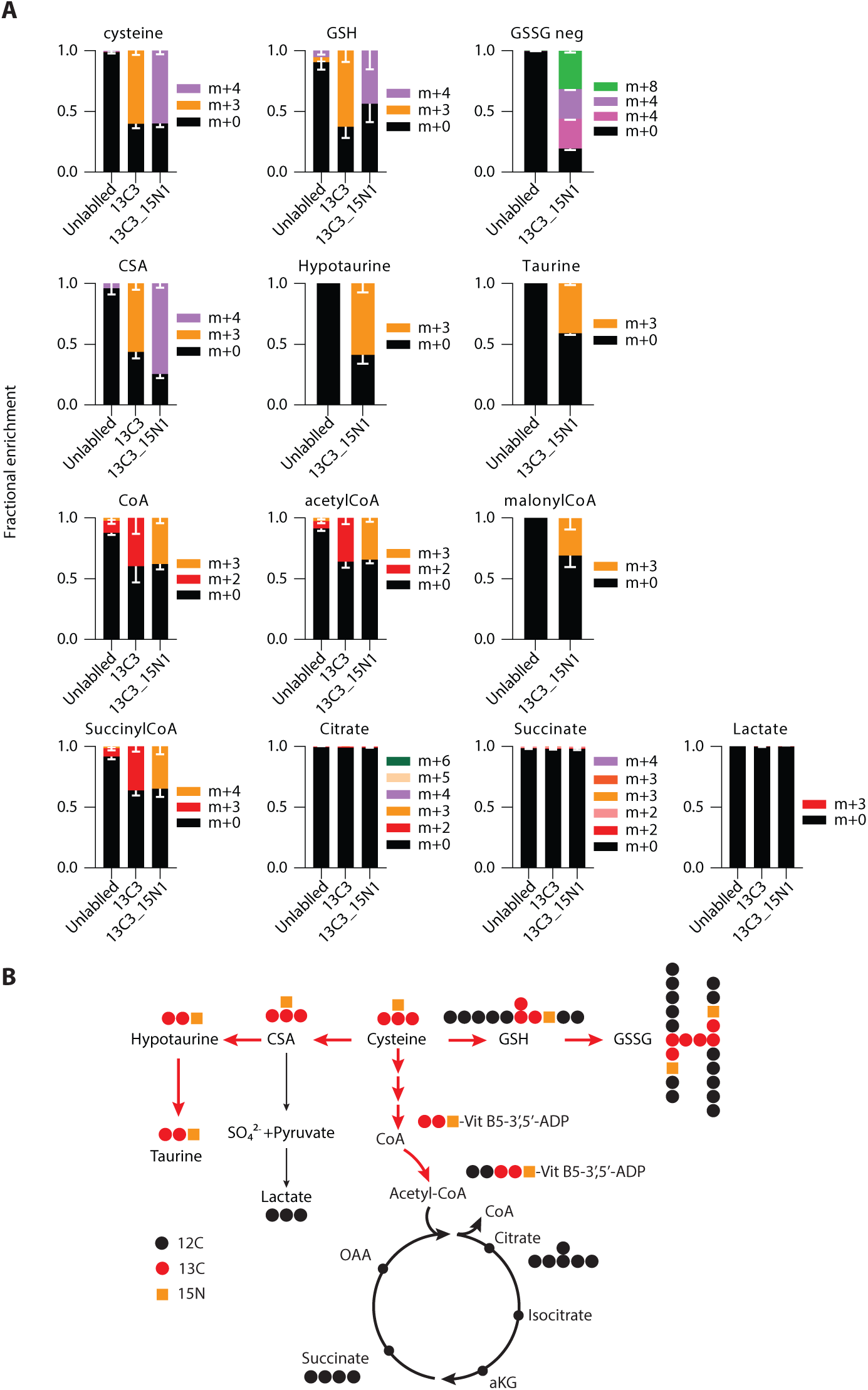
Cysteine fuels *de novo* CoA/acetyl-CoA metabolism upon fasting. A) Mean Fractional enrichment +/- SD of U_^13^C-cysteine (N=10), ^13^C_3__^15^N_1_-cysteine (N=5) or unlabeled samples (N=10) in indicated metabolites measured by LC-MS/MS in whole larvae fast overnight with 5mM tracer. B) Schematic of cysteine metabolism and labelling patterns from U_^13^C-cysteine and ^13^C_3__^15^N_1_-cysteine tracers. Red arrows indicate main cysteine flux.

**Figure 5:**
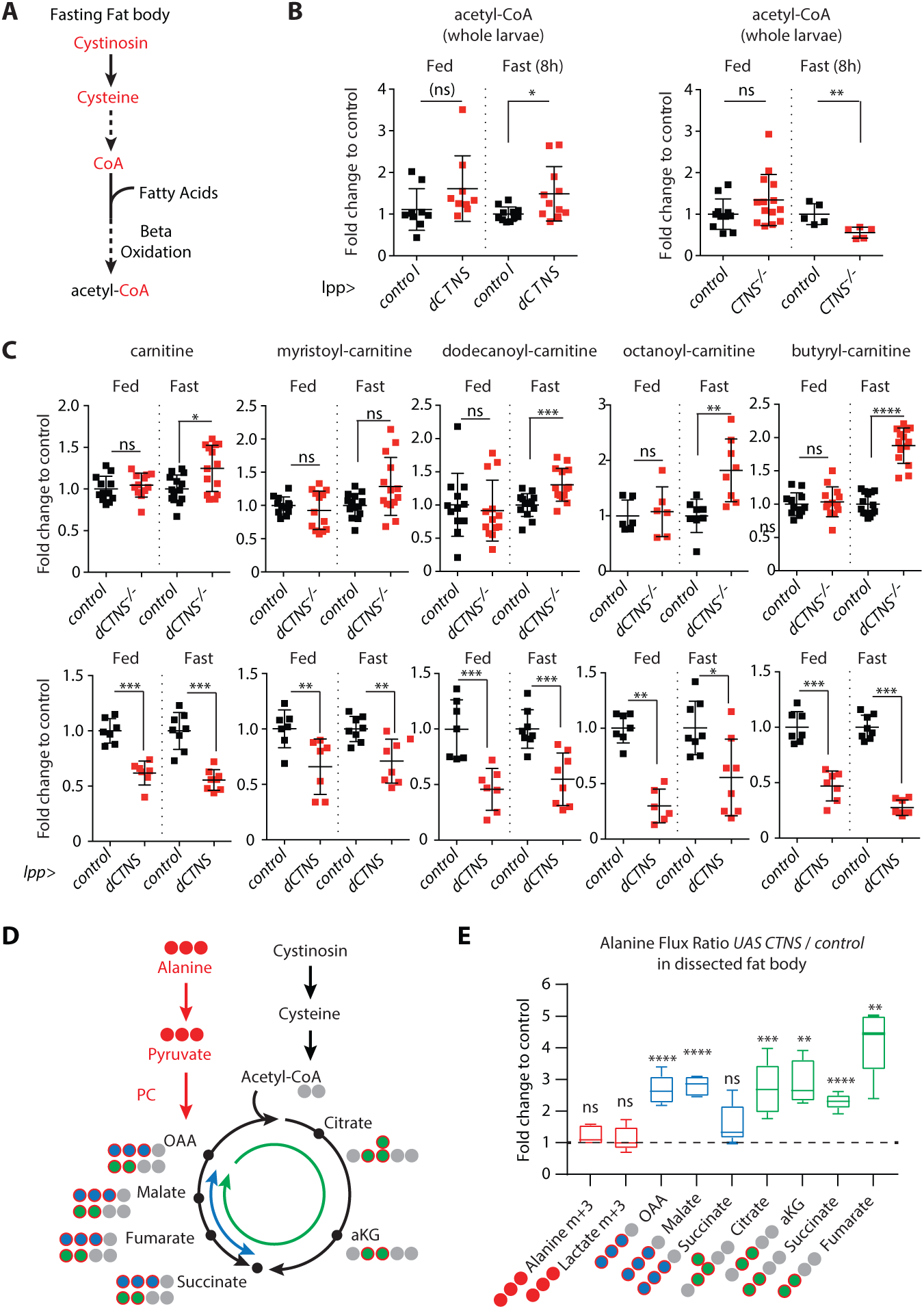
Cysteine metabolism promotes β-oxidation to generate acetyl-CoA and increases carbon flux through PC and the TCA cycle. A) Schematic of acetyl-CoA synthesis. B-C) Metabolite levels in whole 3^rd^ instar larvae *dCTNS*^-/-^ and *dCTNS* overexpression in the fat body. D) Schematic of alanine carbon flux into the TCA cycle upon starvation. E) Alanine flux ratio. Fold change *lpp>CTNS* / control (*lpp>attp40*) for the indicated TCA cycle intermediates isotopomers measure by LC-MS/MS in dissected fat bodies from 3^rd^ instar larvae fed a low protein diet with 25mM U-^13^C-alanine for 6 hours.

Increased CoA synthesis from cysteine could play a central role in enlarging the TCA cycle carbon pool in at least two ways. First, by providing the CoA required to accept increasing amounts of carbon from fatty acids to form acetyl-CoA, and second, by promoting anaplerosis of alternative carbon sources via the acetyl-CoA-dependent allosteric activation of PC (*22*). To test whether the increased production of acetyl-CoA supported by lysosomal cysteine efflux could elevate anaplerosis and TCA cycle intermediates, we analyzed alanine anaplerosis in the TCA cycle following *dCTNS* overexpression. We supplemented a fasted diet with [U-^13^C]alanine so that the tracer had negligible contribution to the total alanine pool (Fig. S5B), and followed the anaplerotic flux of alanine in the TCA cycle in dissected fat body (Fig. 5D). We found that *dCTNS* overexpression strongly increased (>2 fold) alanine anaplerosis as well as oxidative flux in the TCA cycle, through citrate synthase (that consumes OAA and acetyl-CoA to generate citrate, Fig. 5E). Altogether, we propose that during starvation, cysteine recycling and metabolism to acetyl-CoA in the fat body supports anaplerosis through PC and flux through citrate synthase, thereby contributing to the accumulation of TCA cycle intermediates, in particular citrate/ isocitrate.

Finally, we addressed a link between the level of TCA cycle intermediates and TORC1 reactivation. Upon starvation, given the fixed and pre-established level of carbons available, elevation of the level of TCA cycle intermediates may indicate the retention of anaplerotic inputs in the TCA cycle at the expense of their extraction for biosynthesis (i.e. cataplerosis). Because cataplerosis promotes amino acid synthesis (*1*), which in turn affects TORC1 signaling (*6*), we analyzed the effect of lysosomal cysteine recycling on individual amino acid pools. Interestingly, *dCTNS* overexpression in the fat body led to depletion of aspartate and downstream nucleotide precursors (IMP, UMP), as well as, to a lesser extent, asparagine and glutamate (Fig. 6A, S6A, B). Conversely, aspartate and UMP were elevated in *dCTNS*^*-/-*^ animals upon fasting. Aspartate is a cataplerotic product of alanine/oxaloacetate (*1*), a process that involves Got2. Thus, data suggest that cysteine metabolism transiently traps anaplerotic carbons into the TCA cycle upon fasting, away from their immediate extraction for biosynthesis. Further studies will be required to understand the mechanistic basis of this process, which could involve an effect of acetyl-CoA on oxaloacetate flow through citrate synthase towards the TCA cycle vs. Got2 and aspartate synthesis upon fasting. The mild effect on glutamate remains unclear at this point, and may be connected to the metabolism of aspartate as glutamate serves as nitrogen donor for transamination through Got2 during aspartate synthesis. Nevertheless, we analyzed whether cysteine metabolism could regulate growth and antagonize TORC1 reactivation at least through limiting the availability of glutamate and aspartate, because these amino acids are critical regulators of cell growth and have previously been implicated in TORC1 signaling (*6, 23*). To do so, we aimed at replenishing aspartate and glutamate levels in *dCTNS* overexpressing animals using dietary supplementation with anaplerotic amino acids in excess. We found that combinatorial treatments with alanine, aspartate, glutamate and proline (that replenishes glutamate in flies(*24*)) restored normal levels of aspartate and glutamate without compromising *dCTNS*-induced elevation of TCA intermediates (Fig. S6C, D). Strikingly, these treatments rescued the developmental delay induced by *dCTNS* overexpression in the fat body (Fig. 6B). Similarly, co-supplementation of cysteine with excess of single amino acids including alanine, aspartate, glutamate and proline each rescued cysteine-induced growth suppression upon fasting (Fig. 6C). Finally, combinatorial amino acid treatments restored TORC1 activity upon fasting following *dCTNS* overexpression in the fat body (Fig. 6D). Consistent with the importance of aspartate synthesis, loss of Got2 blocked cell growth, induced autophagy and affected TORC1 targets (Fig. S6E), in line with previous reports (*6*). Altogether, we show that cysteine metabolism regulates anaplerotic carbon flow in the TCA cycle and the level of amino acids, in particular aspartate. We propose that cysteine converts the TCA cycle into a reservoir of carbons while limiting their extraction for biosynthesis. This process spares nutrients to survive a starvation period while resetting TORC1 reactivation to a threshold that maintains minimal growth without compromising autophagy.

**Figure 6:**
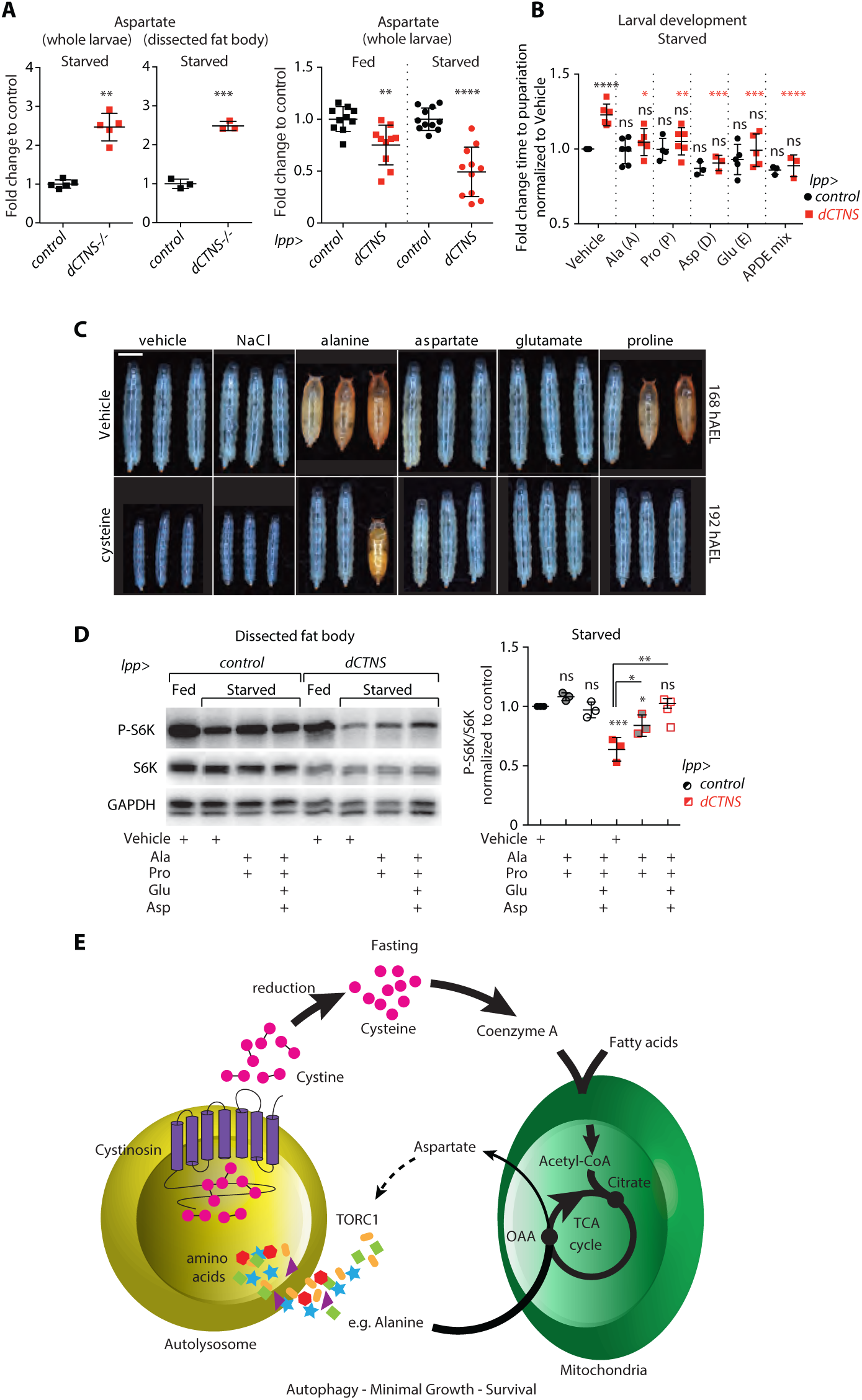
Cysteine metabolism regulates TORC1 and growth through aspartate levels. A) Relative levels of aspartate in 85h AEL larvae *dCTNS*^-/-^ (whole larvae and fat body) and following Cystinosin overexpression in the fat body. B) Amino acid supplementation suppresses the developmental delay induced by *dCTNS* overexpression in the larval fat body. Fold change time to pupariation for larvae fed a minimal diet with or without supplementation with the indicated amino acids (Ala, Pro, Glu: 5 mM, Asp: 10 mM). C) Pictures of aged matched animals fed a minimal diet with or without 5 mM cysteine with or without the indicated metabolites (25 mM each). Scale bar: 1 mm. D) dCTNS-induced TORC1 inhibition is reversed by supplementation with the indicated amino acids. P-S6K levels in fat body from fed and fasted (6h) larvae of indicated genotypes. 70-72h AEL old larvae were transferred to a minimal diet with or without supplementation with the indicated amino acids (Ala, Pro: 20 mM, Asp, Glu: 10 mM). Control is GFP-i. E) Model: cysteine regulates the reactivation of TORC1 upon fasting by limiting biosynthesis from oxaloacetate through the levels of acetylCoA that drives anaplerotic substrates into the TCA cycle. B, D) Mean +/- SD, ^ns^, P≥0.05; *, P≤0.05; **, P≤0.01; ***, P≤0.005; ****p≤0.0001 (Details in Methods).

## Discussion

Maintaining cellular homeostasis upon nutrient shortage is an important challenge for all animals. On the one hand, downregulation of mTORC1 is necessary to limit translation, reduce growth rates and engage autophagy. On the other hand, minimal mTORC1 activity is required to promote lysosomes biogenesis, thus maintaining autophagic degradation necessary for survival (*8*). Here, we show that TORC1 reactivation upon fasting integrates biosynthesis of amino acids from anaplerotic inputs into the control of growth. The regulation of at least aspartate levels appears critical during this process, likely because it serves as precursor for various macromolecules including other amino acids and nucleotides, which in turn impinge on TORC1 activity (*25*). We identify another critical layer of regulation, whereby cysteine recycling via the lysosome fuels acetyl-CoA synthesis and prevents reactivation of TORC1 above a threshold that would compromise autophagy and survival upon fasting. We show that reactivation of TORC1 upon fasting is not passively controlled by the extent of amino acid remobilized from the lysosome. Instead, cysteine metabolism supports an increased incorporation of the carbons from these remobilized amino acids into the TCA cycle. We propose that the remobilized amino acids are transiently stored in the form of TCA cycle intermediates compartmentalized in the mitochondria, thereby regulating their accessibility to maintain autophagy over a starvation period. Interestingly, this pathway is self-regulated by autophagy, as autophagic protein degradation controls cystine availability through the lysosomal CTNS transporter. Thus, in contrast to fed conditions where amino acid transporters at the plasma membrane maintain high cytosolic levels of leucine and arginine that can directly be sensed by members of the TORC1 machinery *(3)*, TORC1 reactivation in prolonged starvation is regulated indirectly by lysosome-mitochondrial crosstalk.

Multiple functions of cysteine impinge on cellular metabolism, including tRNA thiolation, the generation of hydrogen sulfide, the regulation of HIF transcription factors, and its antioxidant function through glutathione synthesis (*26-28*). Here, we show that CoA is a main fate of cysteine that affects oxidative metabolism in the mitochondria, which is source of reactive oxygen species (ROS). Thus, we speculate that the antioxidant function of cysteine could be coupled to its effects on the mitochondria to buffer ROS production. Supplementation with cysteine or modified molecules such as N-acetyl-cysteine (NAC) can be used to efficiently buffer oxidative stress, a process under current investigation to alleviate symptoms of a variety of diseases involving oxidative stress or glutathione deficiency, including cystinosis (*29-32*). In addition, cysteine or NAC extends lifespan in flies, worms and mice, and mice fed NAC show a sudden drop in body weight, similar to dietary restriction (*33, 34*). Our results may challenge the idea that cysteine acts strictly through its antioxidant function since we demonstrate that cysteine supplementation restricts the availability of particular amino acids and limits mTOR activity - processes known to extend lifespan.

In summary, we demonstrate that cysteine metabolism acts in a feedback loop involving *de novo* coenzyme A synthesis, the TCA cycle and amino acid metabolism to limit mTORC1 reactivation upon prolonged fasting and maintain autophagy. We show that this pathway is needed for developing organisms to balance growth and survival during periods of food shortage.

## Acknowledgements

We thank Eric Baehrecke, Christen Mirth, Pierre Léopold, Aurelio Teleman, Frederik Wirtz-Peitz, Yohanns Bellaïche, Isabelle Gaugue, Sylvia Sanquer, Anne-Claire Boschat, Mary A. Lilly, the TRIP (http://www.flyrnai.org/TRiP-HOME.html), BDSC and VDRC stock centers for providing stocks and reagents. We thank Corinne Antignac, Lewis C. Cantley, Bruno Gasnier, Joshua D. Rabinowitz and David M. Sabatini for comments on the manuscript. We thank the Imagine Microscopy platform for assistance with microscopy.

## Author contributions

P.J., Z.M., C.D., M.S. and N.P designed the experiments; P.J., Z.M., A.P., M.D., J.A., and I.N performed the experiments; P.J., Z.M., M.S. and N.P. participated in interpretation of data and P.J wrote the manuscript with inputs from Z.M., C.C.D, M.S. and N.P.

## Data and materials availability

All data is available in the main text or the supplementary materials.

## Supplementary Materials

Materials and Methods

Supplementary Text S1, S2

Figures S1-S6

Tables S1

References (*35-47*)

## References and Notes

1. D. A. Cappel et al., Pyruvate-Carboxylase-Mediated Anaplerosis Promotes Antioxidant Capacity by Sustaining TCA Cycle and Redox Metabolism in Liver. Cell Metab 29, 1291–1305.e1298 (2019).

2. L. R. Gray et al., Hepatic Mitochondrial Pyruvate Carrier 1 Is Required for Efficient Regulation of Gluconeogenesis and Whole-Body Glucose Homeostasis. Cell Metab 22, 669–681 (2015).

3. R. L. Wolfson, D. M. Sabatini, The Dawn of the Age of Amino Acid Sensors for the mTORC1 Pathway. Cell Metab 26, 301–309 (2017).

4. R. C. Scott, O. Schuldiner, T. P. Neufeld, Role and regulation of starvation-induced autophagy in the Drosophila fat body. Dev. Cell 7, 167–178 (2004).

5. T.-C. Lin et al., Autophagy: Resetting glutamine-dependent metabolism and oxygen consumption. Autophagy 8, 1477–1493 (2012).

6. H. W. S. Tan, A. Y. L. Sim, Y. C. Long, Glutamine metabolism regulates autophagy-dependent mTORC1 reactivation during amino acid starvation. Nature Communications 8, 338 (2017).

7. G. A. Wyant et al., NUFIP1 is a ribosome receptor for starvation-induced ribophagy. Science 360, 751–758 (2018).

8. L. Yu et al., Termination of autophagy and reformation of lysosomes regulated by mTOR. Nature 465, 942–946 (2010).

9. J. Colombani et al., A nutrient sensor mechanism controls Drosophila growth. Cell 114, 739–749 (2003).

10. C. Geminard, E. J. Rulifson, P. Leopold, Remote control of insulin secretion by fat cells in Drosophila. Cell Metab 10, 199–207 (2009).

11. Y. Wei, M. A. Lilly, The TORC1 inhibitors Nprl2 and Nprl3 mediate an adaptive response to amino-acid starvation in Drosophila. Cell Death Differ. 21, 1460–1468 (2014).

12. J. Onodera, Y. Ohsumi, Autophagy Is Required for Maintenance of Amino Acid Levels and Protein Synthesis under Nitrogen Starvation. J. Biol. Chem. 280, 31582–31586 (2005).

13. M. Conrad, H. Sato, The oxidative stress-inducible cystine/glutamate antiporter, system x (c) (-): cystine supplier and beyond. Amino Acids 42, 231–246 (2012).

14. H. Kabil, O. Kabil, R. Banerjee, L. G. Harshman, S. D. Pletcher, Increased transsulfuration mediates longevity and dietary restriction in Drosophila. Proc Natl Acad Sci U S A 108, 16831–16836 (2011).

15. M. Town et al., A novel gene encoding an integral membrane protein is mutated in nephropathic cystinosis. Nat. Genet. 18, 319–324 (1998).

16. Z. Andrzejewska et al., Cystinosin is a Component of the Vacuolar H+-ATPase-Ragulator-Rag Complex Controlling Mammalian Target of Rapamycin Complex 1 Signaling. J Am Soc Nephrol 27, 1678–1688 (2016).

17. B. P. Festa et al., Impaired autophagy bridges lysosomal storage disease and epithelial dysfunction in the kidney. Nat Commun 9, 161 (2018).

18. E. A. Ivanova et al., Altered mTOR signalling in nephropathic cystinosis. J Inherit Metab Dis 39, 457–464 (2016).

19. G. Napolitano et al., Impairment of chaperone-mediated autophagy leads to selective lysosomal degradation defects in the lysosomal storage disease cystinosis. EMBO Mol Med 7, 158–174 (2015).

20. J. Zhang et al., Cystinosin, the small GTPase Rab11, and the Rab7 effector RILP regulate intracellular trafficking of the chaperone-mediated autophagy receptor LAMP2A. J. Biol. Chem. 292, 10328–10346 (2017).

21. W. A. Gahl, F. Tietze, J. D. Butler, J. D. Schulman, Cysteamine depletes cystinotic leucocyte granular fractions of cystine by the mechanism of disulphide interchange. Biochem. J. 228, 545–550 (1985).

22. M. St Maurice et al., Domain architecture of pyruvate carboxylase, a biotin-dependent multifunctional enzyme. Science (New York, N.Y.) 317, 1076–1079 (2007).

23. L. B. Sullivan et al., Aspartate is an endogenous metabolic limitation for tumour growth. Nat. Cell Biol. 20, 782–788 (2018).

24. H. Li et al., Drosophila larvae synthesize the putative oncometabolite L-2-hydroxyglutarate during normal developmental growth. Proceedings of the National Academy of Sciences of the United States of America 114, 1353–1358 (2017).

25. G. Hoxhaj et al., The mTORC1 Signaling Network Senses Changes in Cellular Purine Nucleotide Levels. Cell reports 21, 1331–1346 (2017).

26. K. J. Briggs et al., Paracrine Induction of HIF by Glutamate in Breast Cancer: EglN1 Senses Cysteine. Cell 166, 126–139 (2016).

27. C. Hine et al., Endogenous hydrogen sulfide production is essential for dietary restriction benefits. Cell 160, 132–144 (2015).

28. S. Laxman et al., Sulfur amino acids regulate translational capacity and metabolic homeostasis through modulation of tRNA thiolation. Cell 154, 416–429 (2013).

29. M. AlMatar, T. Batool, E. A. Makky, Therapeutic Potential of N-Acetylcysteine for Wound Healing, Acute Bronchiolitis, and Congenital Heart Defects. Curr. Drug Metab. 17, 156–167 (2016).

30. R. Bavarsad Shahripour, M. R. Harrigan, A. V. Alexandrov, N-acetylcysteine (NAC) in neurological disorders: mechanisms of action and therapeutic opportunities. Brain Behav 4, 108–122 (2014).

31. K. Q. de Andrade et al., Oxidative Stress and Inflammation in Hepatic Diseases: Therapeutic Possibilities of N-Acetylcysteine. Int J Mol Sci 16, 30269–30308 (2015).

32. L. Pache de Faria Guimaraes et al., N-acetyl-cysteine is associated to renal function improvement in patients with nephropathic cystinosis. Pediatr Nephrol 29, 1097–1102 (2014).

33. A. A. Parkhitko, P. Jouandin, S. E. Mohr, N. Perrimon, Methionine metabolism and methyltransferases in the regulation of aging and lifespan extension across species. Aging Cell 18, e13034 (2019).

34. M. V. Shaposhnikov et al., Effects of N-acetyl-L-cysteine on lifespan, locomotor activity and stress-resistance of 3 Drosophila species with different lifespans. Aging 10, 2428–2458 (2018).

35. T. Koyama, C. K. Mirth, Growth-Blocking Peptides As Nutrition-Sensitive Signals for Insulin Secretion and Body Size Regulation. PLoS Biol. 14, e1002392 (2016).

36. P. Karpowicz, Y. Zhang, J. B. Hogenesch, P. Emery, N. Perrimon, The circadian clock gates the intestinal stem cell regenerative state. Cell Rep 3, 996–1004 (2013).

37. Andrew R. Bassett, C. Tibbit, Chris P. Ponting, J.-L. Liu, Highly Efficient Targeted Mutagenesis of Drosophila with the CRISPR/Cas9 System. Cell Reports 6, 1178–1179 (2014).

38. J. Bischof et al., A versatile platform for creating a comprehensive UAS-ORFeome library in Drosophila. Development 140, 2434–2442 (2013).

39. W. C. Lee, C. A. Micchelli, Development and characterization of a chemically defined food for Drosophila. PLoS One 8, e67308 (2013).

40. M. D. Piper et al., A holidic medium for Drosophila melanogaster. Nat. Methods 11, 100–105 (2014).

41. A. M. Troen et al., Lifespan modification by glucose and methionine in Drosophila melanogaster fed a chemically defined diet. Age (Dordr) 29, 29–39 (2007).

42. G. M. Mackay, L. Zheng, N. J. F. van den Broek, E. Gottlieb, in Methods Enzymol., C. M. Metallo, Ed. (Academic Press, 2015), vol. 561, pp. 171–196.

43. K. Hahn et al., PP2A regulatory subunit PP2A-B’ counteracts S6K phosphorylation. Cell Metab 11, 438–444 (2010).

44. R. N. Dilger, S. Toue, T. Kimura, R. Sakai, D. H. Baker, Excess dietary L-cysteine, but not L-cystine, is lethal for chicks but not for rats or pigs. J. Nutr. 137, 331–338 (2007).

45. A. Kumar et al., Homocysteine- and cysteine-mediated growth defect is not associated with induction of oxidative stress response genes in yeast. Biochem. J. 396, 61–69 (2006).

46. Y. Nishiuch, M. Sasaki, M. Nakayasu, A. Oikawa, Cytotoxicity of cysteine in culture media. In Vitro 12, 635–638 (1976).

47. M. Tiebe et al., REPTOR and REPTOR-BP Regulate Organismal Metabolism and Transcription Downstream of TORC1. Dev. Cell 33, 272–284 (2015).

