## Supplementary Data for "Lysosomal cystine mobilization shapes the response of mTORC1 and tissue growth to fasting"

Materials and Methods

#### Fly stocks and maintenance

All flies were reared at 25°C and 60% humidity with a 12-h on/off light cycle on standard lab food. N. Perrimon’s standard lab food: 12.7 g/L deactivated yeast, 7.3 g/L soy flour, 53.5 g/L cornmeal, 0.4 % agar, 4.2 g/L malt, 5.6 % corn syrup, 0.3 % propionic acid, 1% tegosept/ethanol. M. Simon’s standard lab food: 18 g/L deactivated yeast, 10 g/L soy flour, 80 g/L cornmeal, 1% agar, 40 g/L malt, 5% corn syrup, 0.3 % propionic acid, 0.2 % 4-hydroxybenzoic acid methyl ester (nipagin)/ethanol. Density was standardized at least one generation before the experiments. For experiments, larvae were reared on freshly made food. *Lpp-gal4* was a gift from Pierre Léopold; *UAS-tsc1, UAS-tsc2* a gift from Christen Mirth(*43*) ; *yw,hs-Flp; mCherry–Atg8a; Act>CD2>GAL4, UAS–nlsGFP/TM6B* a gift from Eric Baehrecke, *hsFlp; act>CD2>Gal4, UAS nlsGFP* a stock from Norbert Perrimon lab(*44*), *yw, hsFlp, Tub-Gal4>UAS-nlsGFP/FM6;;neoFRT82B, TubGal80/TM6,Tb,Hu* a gift from Allison Bardin, *nprl2^1^*  a gift from M. Lilly and *hsFlp; R4-Gal4, UAS-mCherry-Atg8a; FRT82B UAS-GFP/TM6b* a gift from G. Juhász. The following stocks were obtained from BDSC: *UAS-w^RNAi^* (HMS00045), *UAS-w^RNAi^* (HMS00017), *attp40* (#36304), *attp2* (#36303), *UAS-mCherry-nls* (*#*38425), *UAS-Atg1^RNAi^* (HMS02750), *UAS-Atg18a^RNAi^* (JF02898), *UAS-TSC2^RNAi^* (HM04083), *w^1118^*, *UAS-dCTNS^RNAi^* (HMS00213), *UAS-Got2^RNAi^* (HMJ21924)*.* When comparing the effects of RNAi knockdown or protein overexpression through induction of UAS-dsRNA or UAS-cDNA expression, *UAS-w^RNAi^* (HMS00045), *UAS-w^RNAi^* (HMS00017), *attp2* (#36303) and *attp40* (#36304) were used as controls for the TRIP collection (<http://www.flyrnai.org/TRiP-HOME.html>), and UAS-GFP RNAi for *dCTNS* overexpression. *dCTNS* knockout flies were generated with CRISPR/Cas9 technology according to(*46*). Two sgRNA using oligos (one after the ATG start codon in exon 3: GGTGATGTCATGGGAATCGA, and the other before the translation of the first transmembrane domain in exon 4: GGGCAGTACTCGAAATCAGT) were produced by PCR and *in vitro* transcribed into RNA via MEGAscript™ T7 Transcription Kit (ThermoFisher). RNA was injected into *act-Cas9* flies (from Fillip Port/Simon Bullock). F0 flies were crossed to w;; TM3,Sb/TM6,Tb balancer flies and F1 progenies were screened via PCR by uisng oligos flanking the targeted genome region. Any indel difference > 3 bp was visualized in 4% agarose gel in heterozygot F1 progeny. To generate *dCTNS-mKate2* fusion allele, *mKate2* open reading frame was inserted at the C-terminus of *dCTNS* through CRISPR/Cas9 endogenous tagging strategy using vectors kindly provided from Y. Bellaiche (Curie Institut, Paris). In brief, two 1 kb long homology arms (HR1, HR2) of the *dCTNS* gene flanking the sgRNA-guided Cas9 cutting site were cloned into a vector flanking the ATG/STOP-less *mKate2* allele (HR1-linker-mKate2-loxP-mini-white-loxP-linker-HR2). In addition, two vectors for the expression of sgRNA (sgRNA-1: CCACCGTGACCGATGTTCAAAAT, sgRNA-2: CCGAGCGAAGTGACGACTGAGAA) targeting the C-terminal coding region of *dCTNS* were generated. All three vectors were injected into *vas-Cas9* flies (BDSC#55821) embryos by Bestgene. Progenies were screened for the red eyes (selection marker mini-*white*) and crossed to Cre-expressing flies to remove the mini-*white* by *loxP*/Cre excision. For overexpression of *dCTNS*, *dCTNS* cDNA was cloned into Gateway destination vector pUASg-HA.attB (GeneBank: KC896837) according to(*47*). The plasmid was injected by Bestgene into FlyC31 embryos (BDSC#24482) for *ϕ31*-mediated recombination at a *attP* insertion site on the second chromosome.

Fly food and starvation protocols

Heavy isotope tracers were from Cambridge Isotope Laboratories. All other amino acids and compounds used were from Sigma. In N. Perrimon’s lab compounds in solution were added to the following food mix: 60 g/L sucrose, deactivated yeast as a source of total protein (2 g/L for fast, 4 g/L for mildly fast, 20g/L for fed fly food); 80 g/L cornmeal, 0.35% Bacto Agar, 0.3% propionic acid, 1% tegosept (100g/L in ethanol). Glucose tracing was done without sucrose, and alanine tracing without yeast. In the Simon’s lab fasting food had to be adapted to 6 g/L of deactivated yeast to match control fast (2 g/L) developmental rates observed in the Perrimon lab. For Simon’s lab fed food, the standard lab food was used (see protocol above). In Extended Data Fig. 2 specifically, 100% amino acid is 17g/L deactivated yeast. For starvation on PBS, larvae were placed on paper wipes soaked in PBS in a petri dish.

#### Generation of clones

For autophagy experiments, clones were generated by crossing *yw,hs-flp; mCherry–Atg8a; Act>CD2>GAL4, UAS–nlsGFP/TM6B* with the indicted UAS lines. Progeny of the relevant genotype was reared at 25°C and spontaneous clones were generated in the fat body due to the leakiness of the heat-shock flipase (*hs-flp*). For *dCTNS*^-/-^ clones, autophagy was analyzed by crossing *w;; neoFRT82B, dCTNS*^-/-^ to *hs-flp; R4-Gal4, UAS-mCherry-Atg8a; FRT82B UAS-GFP/TM6b*. For P-4EBP1 experiments, clones were either generated by crossing *hs-flp; act>CD2>Gal4, UAS nlsGFP* with the indicted UAS lines or, for *dCTNS*^-/-^ clones, by crossing *w;;neoFRT82B, dCTNS*^-/-^ to *yw, hs-flp, tub-Gal4>UAS-nlsGFP/FM6;;neoFRT82B, tubGal80/TM6,Tb,Hu*. F1 embryos collected overnight were heat shocked for 2h at 37°C the following morning.

### Amino acid screen

We fed larvae a diet with reduced yeast extract/proteins (50% of normal diet), systematically added individual amino acids to the food, and monitored the time to pupariation as a proxy for growth rate. The following mix was diluted 1:1 with amino acids solutions in water: 10g/L Agar; 120g/L Sucrose; 17g/L Deactivated Yeast extract; 80g/L Cornmeal; 6ml/L Propionic Acid; 20ml/L Tegosept. The amount of amino acids added to the food was determined based on those used in tissue culture growth supplements (See Table S1)(*40-42*).

Food intake

Larvae were synchronized in L1 and reared on the indicated food types until mid-2^nd^ instar. Larvae were then transferred on the same food type supplemented with 0.5% weight/volume erioglaucine disodium salt (Sigma) for 2 hours. Samples were homogenized in 200 µl of PBS and absorbance of the dye in the supernatant was measured at 625 nm. Results were normalized to protein content.

#### Developmental timing

Three-day-old crosses were used for 3-4 hour periods of egg collection on standard lab food. Newly hatched L1 larvae were collected 24 hours later for synchronized growth using the indicated diets at a density of 30 animals/vial. The time to develop was monitored by counting the number of animals that underwent pupariation, every two hours in fed conditions or once/twice a day in starved conditions. The time at which half the animals had undergone pupariation is reported.

Life span experiments

To generate age-synchronized adult flies, larvae were raised on lab food at low density, transferred to fresh food upon emerging as adults and mated 48h. Animal were anaesthetized with low levels of CO_2_ and males sorted at a density of ten per vial. Each condition contained 8-10 vials. Each experiment was repeated at least 3 times and the average values of each experiment were used for statistical analysis. Flies were transferred to fresh vials three times per week at which point deaths were scored. After ten days, deaths were scored every day.

Growth curves/pupal weight

Synchronized, newly-hatched L1 larvae were immediately weighed or placed on the indicated food at a density of 30-50 animals/vial. Pools of 20-80 animals were weighed every 24 hours using an analytical scale (Mettler Toledo) and the weight/animal was reported +/- SEM. For pupal weight, two-day-old pupae from vials at a density of 30 animals were weighed in batches of 5-10 pupae. The weights of different batches of larvae from the same vials were averaged and counted as N=1.

Metabolite profiling

For whole body metabolic profiling, 25-38 mid-second instar or 8-15 mid-third instar larvae per sample were collected, snap-frozen in liquid nitrogen and stored at -80°C in extraction buffer (4-6 biological replicates/experiment). For fat body metabolic profiling, fat bodies from 35-40 larvae 96h AEL old were dissected in 20 ul PBS, diluted in 300 ul cold extraction buffer and snap frozen. Tissues were homogenized in extraction buffer using 1 mm zirconium beads (Next Advance, ZROB10) in a Bullet Blender tissue homogenizer (Model BBX24, Next Advance). Metabolites were extracted using 80 % (v/v) aqueous methanol (x2 sequential extractions with 300-600 ul) and metabolites pelleted by vacuum centrifugation. Pellets were resuspended in 20 ul HPLC-grade water and metabolomics data were acquired using targeted liquid chromatography tandem mass spectrometry (LC-MS/MS). A 5500 QTRAP hybrid triple quadrupole mass spectrometer (AB/SCIEX) coupled to a Prominence UFLC high-performance LC (HPLC) system (Shimadzu) was used for steady-state analyses of the samples. Selected reaction monitoring (SRM) of 287 polar metabolites using positive/negative switching with hydrophilic interaction LC (HILIC) was performed. Peak areas from the total ion current for each metabolite SRM Q1/Q3 transition were integrated using MultiQuant version 2.1 software (AB/SCIEX). The resulting raw data from the MultiQuant software were normalized by sample weight for whole animal. Fat body samples were normalized by the mean protein content measured from duplicate dissection for each condition. Data were analyzed using Prism informatic software. Alternatively, collected larvae were rinsed with water, 70% ethanol and PBS to remove food and bacteria, snap-frozen in liquid nitrogen and stored at -80°C until extraction in 50% methanol, 30% ACN, and 20% water. The volume of extraction solution added was adjusted to larvae mass (40mg/ml), samples were vortexed for 5 min at 4°C, and then centrifuged at 16,000 g for 15 minutes at 4°C. Supernatants were collected and analyzed by LC-MS using a QExactive Plus Orbitrap mass spectrometer equipped with an Ion Max source and a HESI II probe and coupled to a Dionex UltiMate 3000 UPLC system (Thermo, USA). An SeQuant ZIC-pHilic column (Millipore) was used for liquid chromatography separation(*34*). The aqueous mobile-phase solvent was 20 mM ammonium carbonate plus 0.1% ammonium hydroxide solution and the organic mobile phase was acetonitrile. The metabolites were separated over a linear gradient from 80% organic to 80% aqueous for 15 min and detected across a mass range of 75–1,000 m/z at a resolution of 35,000 (at 200 m/z) with electrospray ionization and polarity switching mode. Lock masses were used to insure mass accuracy below 5 ppm. The peak areas of different metabolites were determined using TraceFinder software (Thermo) using the exact mass of the singly charged ion and known retention time on the HPLC column. In total, each metabolic profiling experiment was performed at least two times with 3-7 biological replicates per genotype.

#### TCA cycle isotopomer method from U-^13^C-cysteine, ^13^C_3_-^15^N_1_-cysteine, U-^13^C-glucose and U-^13^C-alanine

Fed and starved mid-third instar animals were supplemented with the indicated concentrations of U-^13^C-cysteine, ^13^C_3__^15^N_1_-cysteine, U-^13^C-glucose, U-^13^C-alanine or vehicle in the food during the indicated time. For tracing experiments on low protein diet (starved), animals were pre-starved 1 hour on PBS before being transferred to the relevant tracer/food. Sample were collected (≥5 biological replicates for labelled conditions, ≥4 biological replicates for unlabeled condition), and intracellular metabolites were extracted using 80% (v/v) aqueous methanol. Q1/Q3 SRM transitions for incorporation of 13C labeled metabolites were established for polar metabolites isotopomers and data were acquired by LC-MS/MS. Peak areas were generated using MultiQuant 2.1 software. Unless otherwise indicated, the mean peak areas from unlabeled conditions were used for background determination and were subtracted from each corresponding ^13^C labelled dataset.

Cysteine measurement (other than LC-MS/MS)

25-40 mid-second instar animals were homogenized in cold PBS 0.1% Triton and centrifuged at 4°C. Cysteine measurement was performed in triplicate from the supernatant using the MicroMolar Cysteine Assay Kit (ProFoldin, CYS200) according to the manufacturer’s instructions. Data were normalized to protein content.

Cystine measurements

Larvae were washed three time (water, shortly in 70% ethanol, and finally in PBS), dried on tissue paper and 10 larvae/sample were shock frozen in liquid nitrogen and stored at -80°C until lysis. Larvae were lysed in 80 μl of 5.2 mM N-ethylmaleimide, centrifuged 10 min at 4°C and 75 μl supernatant was deproteinized by addition of 25 μl of 12% sulfosalicylic acid. Protein-free supernatants were kept frozen at -80°C until analysis and centrifuged at 1,200 x g before use. Cystine quantification was performed using AccQ-Tag Ultra kit (Waters®) on a UPLC-xevoTQD system (Waters) according to the manufacturer’s recommendations. Ten µL of samples were mixed with 10 µL of a 30 µM internal standard solution (stable isotope of cystine), 70 µL of borate buffer and 20 µL of derivative solution and incubated at 55°C for at least 10 minutes. Derivatized samples were diluted with 150 µL of ultrapurified water and 5 µL of the final mix were injected in the triple quadrupole mass spectrometer in positive mode. Transitions used for derivatized cystine quantification and the internal standard were 291.2>171.1 and 294.2>171.1, respectively. Cystine values were normalized by protein content using the Lowry’s method on protein pellets.

#### Immunostaining

Tissues from 68-85 hours AEL larvae were dissected in phosphate-buffered saline (PBS) 2% formaldehyde at room temperature, fixed 20-30 min in 4% formaldehyde, washed twice 10 min in PBS 0.3% Triton (PBST), blocked 30 min (PBST, 5% BSA, 2% FBS, 0.02% NaN3), incubated with primary antibodies in the blocking buffer overnight and washed 4 times for 15 min. Secondary antibodies diluted 1:200 or 1:500 in PBST were added for 1 hour and tissues washed 4 times before mounting in Vectashield/DAPI. Rabbit anti-P-4EBP1 was from Cell Signaling Technologies (CST 236B4, #2855) and diluted 1:500, rabbit anti-tRFP was from Evrogen (#AB233) and used against mKate2 to stain cystinosin-mKate2. Samples were imaged using Zeiss LSM 780 and Leica TCS SP8 SMD confocal systems with a 40x water or 40x oil immersion objective and images were processed with Fiji software.

#### Western Blots

Tissues from 10-30 animals were dissected in CST lysis buffer (Cat#9803) containing 2x protease inhibitor (Roche, 04693159001) and 3x phosphatase inhibitor (Roche, 04906845001), and homogenized using 1 mm zirconium beads (Next Advance, ZROB10) in a Bullet Blender tissue homogenizer (Model BBX24, Next Advance). Protein content was measured to normalize samples, 2x Laemmli Sample Buffer (Biorad) was added and samples boiled 6 min @ 95°C. Lysates were resolved by electrophoresis (Mini-PROTEAN TGX Precast Gels, BioRad, PAGEr EX Gels, Lonza, or home-made SDS gels), proteins transferred onto PVDF membranes (Immobilon P, Millipore), blocked in Tris-buffered saline with or without 0.1% Tween-20 buffer containing 3-6% BSA or 5% milk, and probed with P-S6K antibody (1:1000, CST 9209). After P-S6K was revealed, membranes were stripped for 5-30 min (Restore PLUS Buffer, Thermo Scientific #46430), washed, blocked in PBS Tween-20 buffer containing 5% dry milk, and probed with S6K antibody (1:10000, a gift from Aurelio Teleman(*35*)). For normalization blots were probed with GADPH antibody (1:5000, GeneTex GTX100118). Data show representative results from at least 2 or 3 biological replicates (see quantification plots). Horseradish peroxidase (HRP) conjugated secondary IgG antibodies (1:10000) were used together with the SuperSignal West Dura Extended Duration Substrate (Thermo Scientific #34076) to detect the protein bands.

Statistics

Experiments are presented with whisker plots or the mean +/- SD or SEM. P values and significance: ^ns^, P≥0.05; *, P≤0.05; **, P≤0.01; ***, P≤0.005; ****, P≤0.0001. **Life span experiments:** Fig. 3A: N=2; Fig. 3C, E, G (N≥4): significance was determined by a two-tailed t-test (Mann-Whitney). N=1 means average of 8-10 vials per genotype and condition in 1 experiment. **Larval development (Pupariation assay):** Fig. 3B and Fig. S3G: significance was determined by a two-tailed t-test (Mann-Whitney); Fig. 3D, F, and Fig. 6B: significance was determined by one-way ANOVA followed by a Bonferroni multiple comparisons test. **Cysteine measurements (Profoldin kit):** Fig. S3A, E, F: significance was determined by a two-tailed t-test (Mann-Whitney). **Western blots:** Fig 1A: spline curve represent the trend over multiple experiments. Fig. 1D, Fig. S1A: 2 ways ANOVA followed by Sidak’s multiple comparisons test, Fig. 2D, F, and Fig. 6D: significance was determined by one-way ANOVA followed by a Bonferroni multiple comparisons test. **Untargeted/targeted metabolomics:** Fig. 1B plots left, Fig. S6C: significance was determined by one-way ANOVA followed by a Tukey’s multiple comparisons test. Fig. 1B plots right, Fig. 2H, Fig. 5B, C, E, Fig. 6A, Fig. S1D, E, Fig. S2G, Fig. S5A, B, Fig., S6A, B, D, Fig. S3D: significance was determined by a two-tailed t-test (Mann-Whitney). **Pupal weight**: Fig. S2C: significance was determined by unpaired two-tailed t-test. **Food intake**: Fig. S2D: significance was determined by unpaired two-tailed t-test.

Supplementary Text

#### Supplementary Text 1: An amino acid supplementation screen reveals cysteine as a growth suppressor

To evaluate the effects of individual amino acids on growth, we developed an amino acid supplementation screen on developing *Drosophila* larvae and measured their development rate (Supplementary Materials). The screen identified cysteine as a strong growth suppressor (Fig. 2A), an effect that could be due to the cytotoxicity of cysteine previously reported in cell culture, yeast, and chicks(*37-39*). However, we found that the effect of cysteine supplementation was diet-dependent, with cysteine strongly suppressing growth upon starvation while having weaker effect in fed animals (Extended Data Fig. 2A-D), mitigating toxicity as a unique explanation for this result. In addition, although the effect of cysteine on growth was dose-dependent (Extended Data Fig. 2A), variation in cysteine intake between fed and starved conditions was not sufficient to explain the diet-dependent toxicity (Extended Data Fig. 2D). Therefore, we conclude that the growth-suppressive effect of cysteine was multifactorial and decided to analyze the endogenous role of intracellular cysteine.

#### Supplementary Text 2: Developmental delay versus starvation sensitivity

Gain and loss of function of *dCTNS* have opposite effect on TORC1 but showed similar phenotype in term of larval development: an increase in the time to pupariation. To avoid confusion between opposite processes that lead to similar phenotypes in appearance, we adapted our nomenclature accordingly. Because the loss of *dCTNS* caused reduced cellular cysteine, upregulation of TORC1, inhibition of autophagy, and that cysteine and rapamycin treatments rescued/accelerated the time to pupariation (Fig. 3), we termed *dCTNS^-/-^* developmental phenotypes “starvation sensitivity”. Accordingly, *dCTNS^-/-^* or depletion of dCTNS in fat body did not affect development of fed larvae. By contrast, *dCTNS* overexpression increased cellular cysteine, downregulated TORC1 and induced autophagy. In agreement with TORC1 loss of function delaying larval growth and development, we termed the developmental phenotype of *dCTNS* overexpression “developmental delay”. Consistently, *dCTNS* overexpression retarded development in both fed and starved conditions.

**Figure Legend**

**Fig. S1: Autophagy-dependent TORC1 reactivation and functional TCA cycle upon fasting.** A, B) TORC1 reactivation upon fasting requires autophagy. P-S6K levels in dissected fat body from control and fat body-expressing *Atg1 RNAi* and *Atg18a RNAi* larvae starved for the indicated time. C) ^13^C fractional enrichment (%) for metabolite isotopologues (different number of ^13^C atoms) and isotopomers (^13^C at different atomic position) measured in whole larvae. Starved animals were supplemented for 4 hours with 25 mM U-^13^C_6_-glucose. OAA, oxaloacetate. D) Relative NADH/NAD+ measured by LC-MS/MS in fed and starved conditions for control animals. E) Fold change in alanine levels in fed and starved (6h on PBS) condition following PC knowdown (whole larvae).

**Fig. S2: Cysteine suppresses growth in a diet-dependent manner and increases TCA cycle intermediate levels.** A) Cysteine suppresses growth in a dose and diet dependent manner. Mean fold change time to pupariation (*w^1118^* larvae) as a function of total protein content and cysteine concentration supplemented in the food. Data are normalized to control for each diet. B) Cysteine supplementation affects larval growth all along development. Growth curves (mean larval weight (mg) as a function of time (hours AEL)) for *w1118* larvae fed *ad libitum* a 100% [AA] (top) or 10% [AA] (bottom) diet with 5 mM cysteine or vehicle. N=3. Pictures show age-matched larvae at 96 and 144 hours AEL in fed and fast conditions, respectively. Scale bar 1mm. C) Animals fed cysteine are reduced in adult size and this process is diet dependent. Pupal weight +/- SEM (mg) of *w^1118^* larvae fed *ad libitum* 300, 100, 50 and 20% total protein diet with 10 mM cysteine or vehicle. D) Diet affects food intake whereas cysteine supplementation does not. Food intake normalized +/- SD to protein content of larvae raised on the indicated diet with 5 mM cysteine or vehicle. E) Cysteine treatment inhibits TORC1 activity. P-S6K levels in dissected fat body from larvae fed a full diet with the indicated concentrations of cysteine or vehicle for the indicated time. Prolonged cysteine treatment (bottom panel) cause downregulation of S6K levels. F) Constitutive TORC1 activity partially suppresses the effect of cysteine on growth. Fed and fasted, control and Gator1 mutant (*nprl2*) larvae supplemented with 5 mM cysteine or vehicle *ad libitum*. G) Cysteine increases the level of TCA cycle intermediates. Relative metabolites levels +/- SD measured by LC-MS/MS in whole mid-second instar *w^1118^* larvae fed the indicated diet *ad libitum* (fast, fasted; mild fast, mildly fasted), with 10 mM cysteine or vehicle. c, d, g) ^ns^, P≥0.05; **, P≤0.01, ***, P≤0.005; ****p≤0.0001 (for further details about statistics see Methods).

**Fig. S3: Lysosomal cystine efflux through *dCTNS* regulates TORC1 reactivation and autophagy.** A) Autophagy controls cysteine levels upon fasting. Relative cysteine levels measured from whole larvae. GFP-i, control background. B) Schematic of the *dCTNS* locus and alleles generated by CRISPR/Cas9. *dCTNS-mKate2*, C-terminal *mKate2* insertion at the endogenous locus; *dCTNS^-/-^*, frameshift mutant causing a premature stop-codon. C) Cystinosin is targeted to the lysosomal membrane. Co-localization of Cystinosin-mKate2 with lysosomal Lamp1-GFP in the fat body. Scale bar, 20 μm and 2 μm. D) *dCTNS^-/-^* larvae accumulate cystine. Relative cystine levels in whole third instar larvae fed a standard diet. E) Cystinosin controls cysteine levels upon fasting. Reative cysteine levels in whole larvae. Controls are heterozygote animals (*dCTNS*^+/-^). F) *dCTNS* overexpression in larval fat body (*lpp*>*dCTNS*) increases cysteine levels. Whole body cysteine levels from fed larvae. Control is GFP-i. G) *dCTNS* overexpression in larval fat body causes a developmental delay. Fold change time to pupariation (Hours AEL). H) Cystinosin limits TORC1 reactivation upon fasting. *dCTNS^-/-^* fat body clones (non-GFP, outlined) and Rheb and TSC1, TCS2 overexpression clones (GFP, outlined) in 80h AEL animals starved for 24h and stained for P-4EBP. DAPI, blue; GFP, green; P-4EBP, red or white. Scale bar 10 μm. I) The TORC1 inhibitor rapamycin restores autophagy in starved dCTNS^-/-^. Fat body *dCTNS^-/-^* clones (non-GFP, outlined) in 80h AEL larvae with fat body-specific expression of mCherry-Atg8a starved for 8h. DAPI, blue; GFP, green; mCherry-Atg8a, red or white. Scale bar 10 μm. a, d-g) Mean +/- SD, **, P≤0.01 (details in Methods).

**Fig. S4: Cysteine metabolism fuels Acetyl-CoA synthesis.** A, B) ^13^C fractional enrichment (%) for metabolite isotopologues and isotopomers measured in whole larvae (A) or dissected fat bodies (B). Starved animals were supplemented for the indicated time with 5 mM ^13^C_3_-cysteine in a fasted diet. Mean +/- SD

**Fig. S5: CTNS affects the levels of fatty acids.** A) Whole body fatty acids levels in 3^rd^ instar *dCTNS*^-/-^ animals and following Cystinosin overexpression in the fat body. B) ^13^C fractional enrichment in alanine in starved flies fed 25mM [U-^13^C]alanine for 6 hours.

**Fig. S6: Cysteine metabolism regulates growth through aspartate levels during starvation.** A, B) Whole body metabolite levels (inosine monophosphate, IMP; uridine monophosphate, UMP) in 3^rd^ instar *dCTNS*^-/-^ animals or following Cystinosin overexpression in the fat body. C) Relative levels of aspartate and glutamate in 85h AEL larvae fed a minimal diet all along development, with or without supplementation with the indicates amino acid cocktails for 8h before collection (Ala, Pro: 20 mM, Asp, Glu: 10 mM). Controls are GFP-i. D) Relative levels of TCA cycle intermediates in 85h AEL larvae fed a minimal diet with or without supplementation with amino acid cocktails for 8h (amino acid concentrations: Ala, Pro: 20 mM, Asp, Glu: 10 mM). Controls are GFP-i. E) Fat body flip-out clones of *Got2-RNAi* (GFP or RFP, outlined) in larvae fed or fasted for 6 hours stained with GFP or P-4E-BP (independent experiments and confocal settings). Atg8a, mCherryATG8a. Unk-GFP is a reporter induced by TORC1 inhibition (*36*). Scale bar 20 μm. A-D) Mean +/- SD; ^ns^, P≥0.05; *, P≤0.05; **, P≤0.01; ***, P≤0.005; ****, P≤0.0001 (Details in Methods).

**Table S1: Amino acid concentrations used in the screen. C**omparative table of concentration of each amino acid (mM) used in Minimum Essential Media (MEM) amino acid supplementation for cell culture, and in our amino acid add-back screen (Fig.2A).
