## Supplemental Figures for "Lysosomal cystine mobilization shapes the response of mTORC1 and tissue growth to fasting"

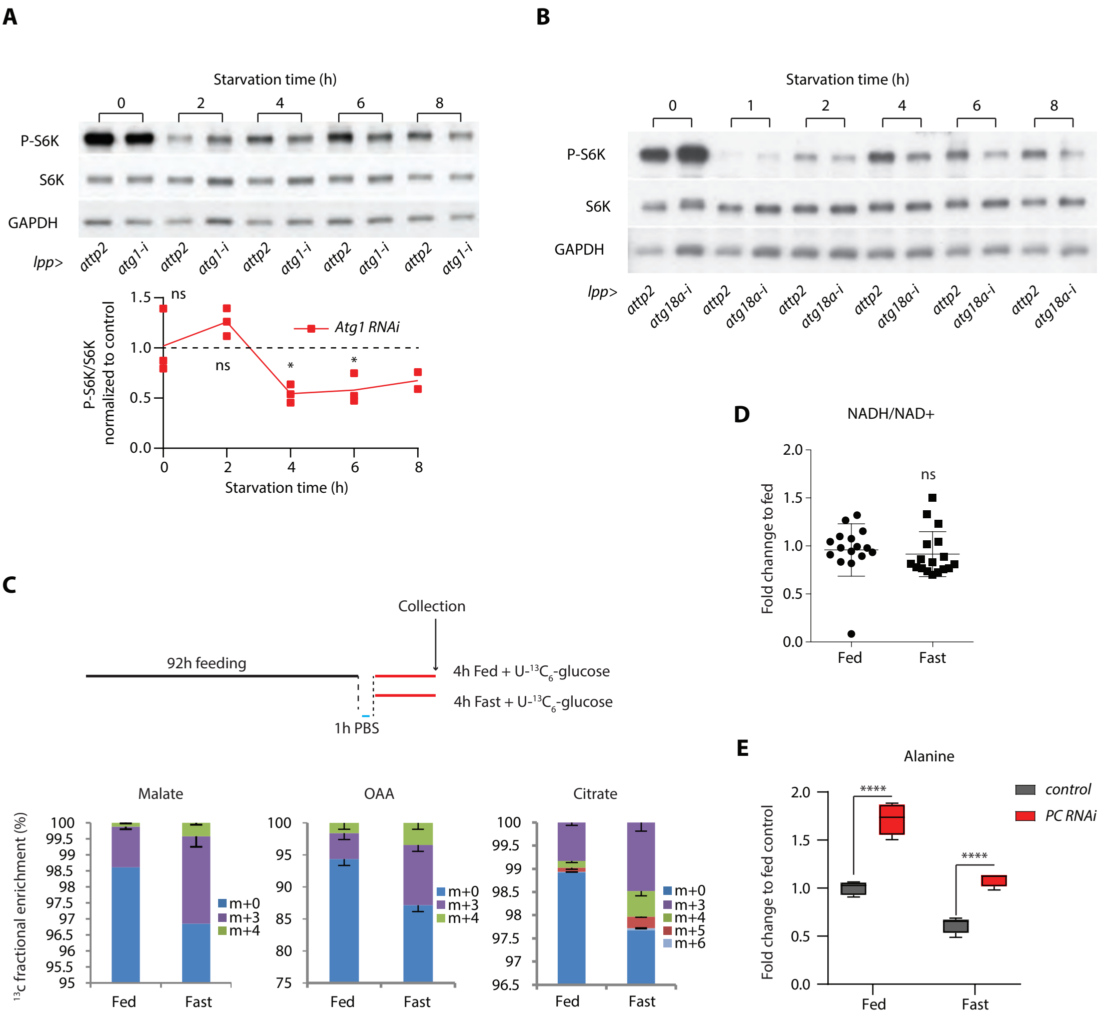


Jouandin et al Fig. S1

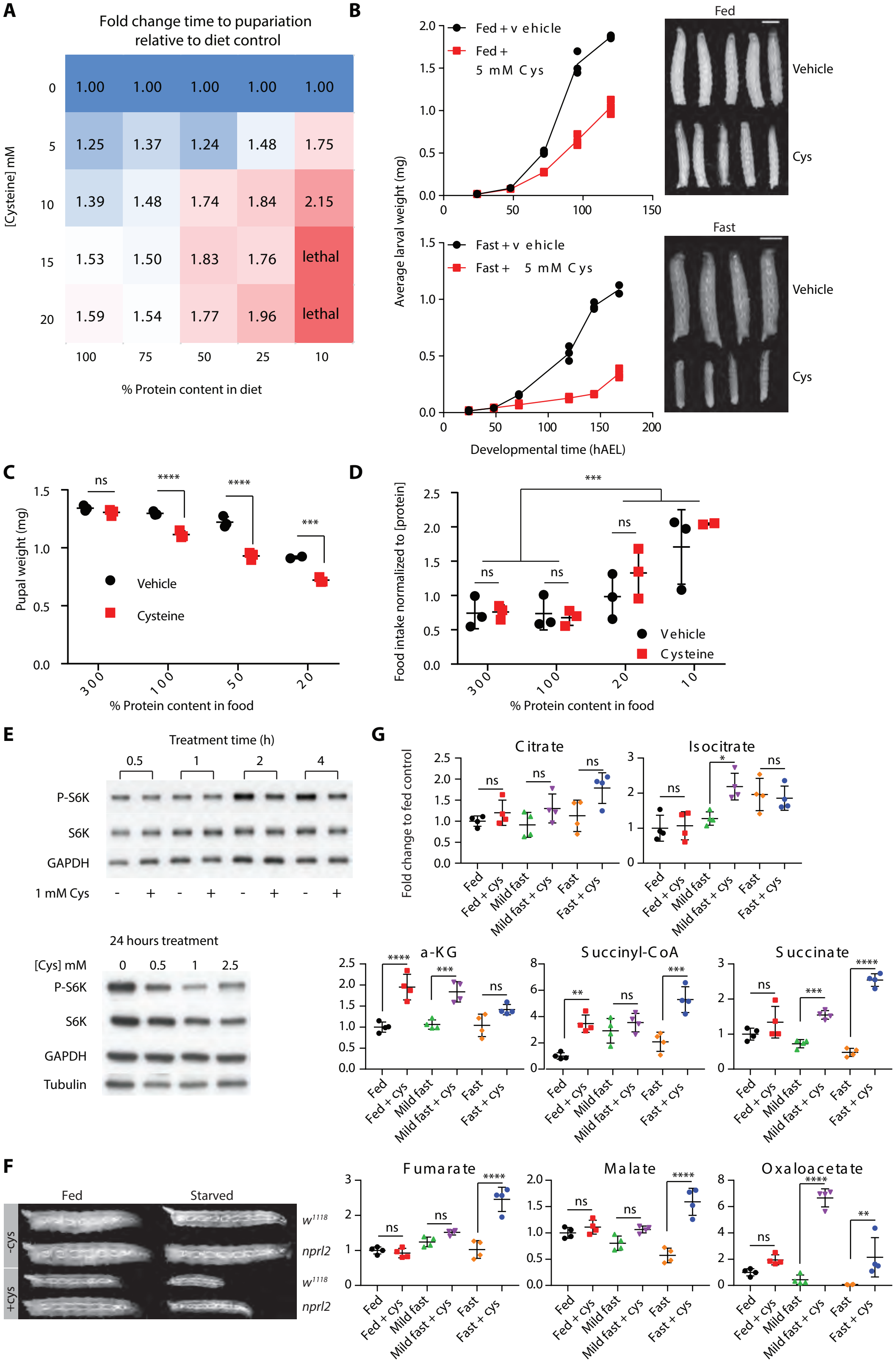


Jouandin et al Fig. S2

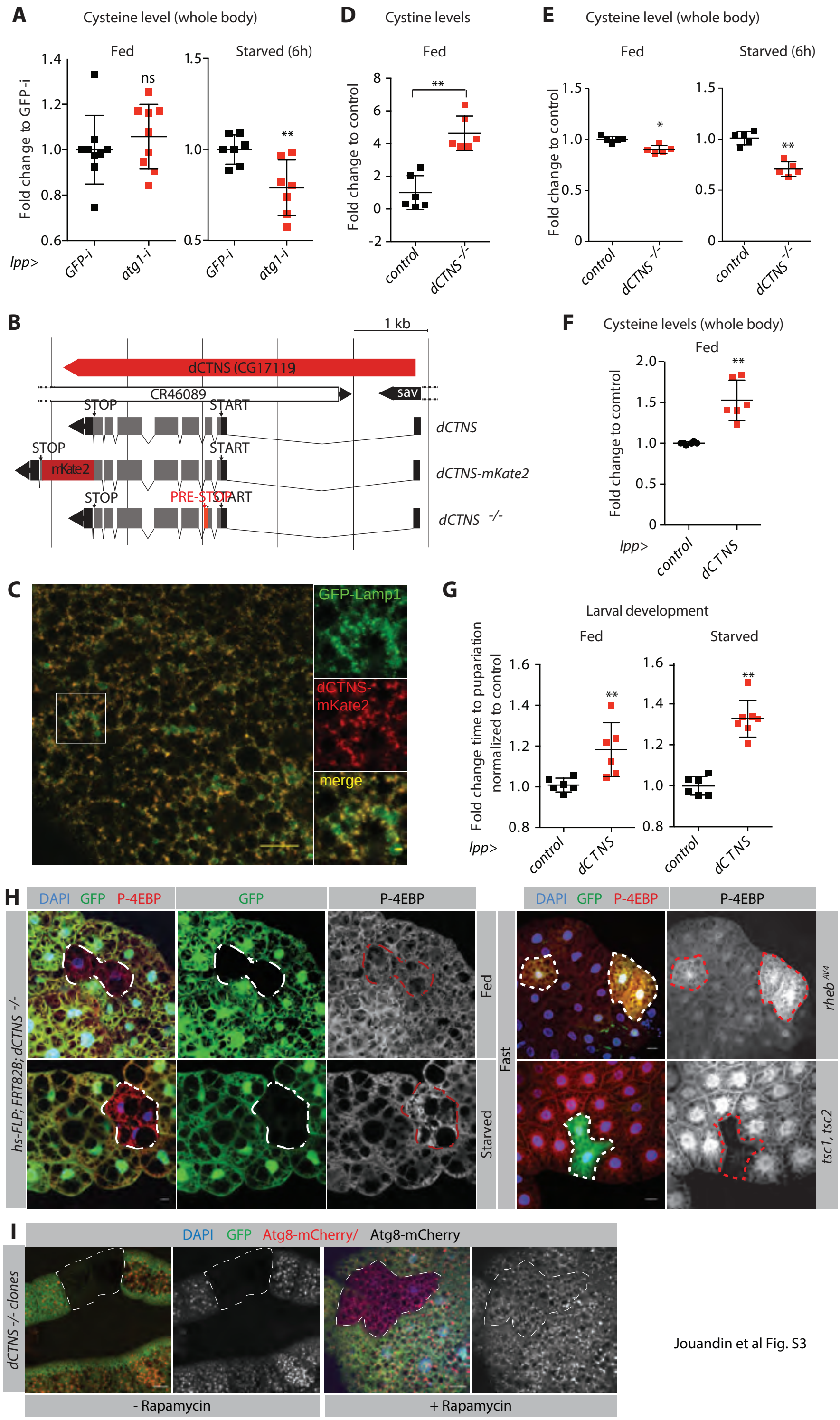


Jouandin et al Fig. S3

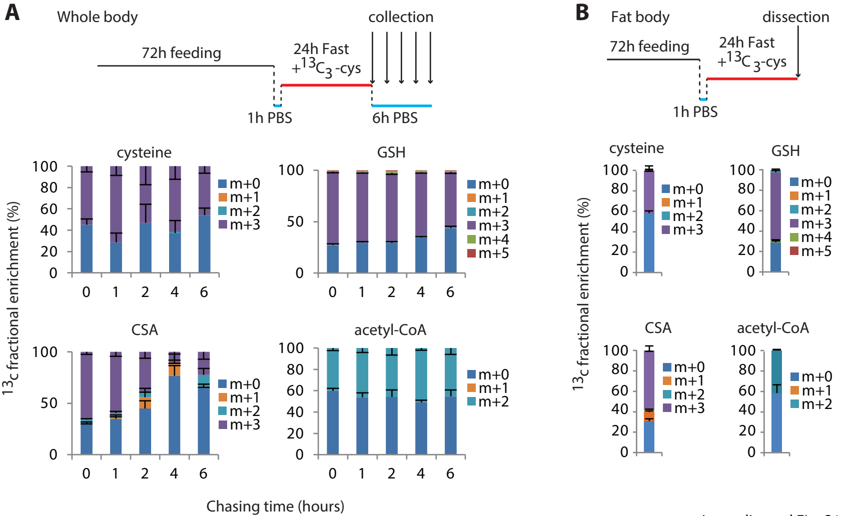


Jouandin et al Fig. S4

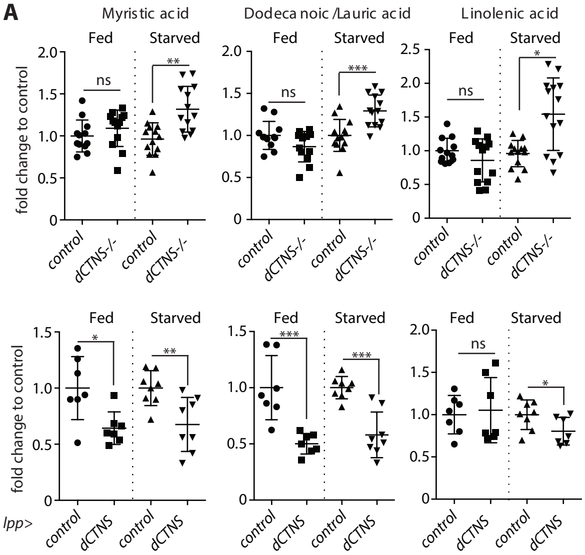


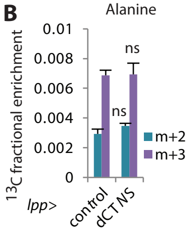


Jouandin et al Fig. S5

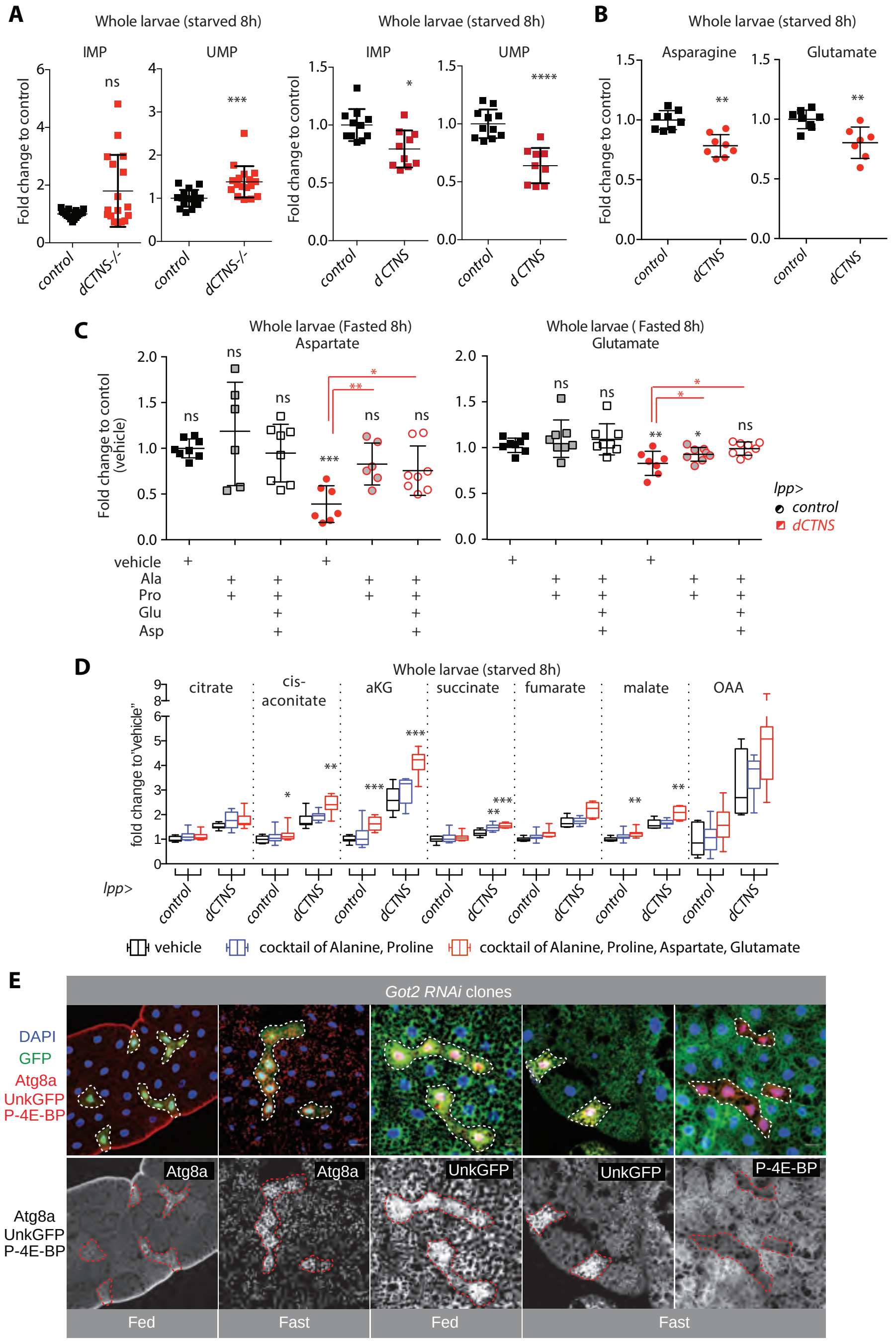


Jouandin et al fig. S6
